# Simultaneous Spectral Differentiation of Multiple Fluorophores in Super-resolution Imaging Using a Glass Phase Plate

**DOI:** 10.1101/2022.07.11.499581

**Authors:** Sanduni I. Fernando, Jason T. Martineau, Robert J. Hobson, Thien N. Vu, Brian Baker, Brian D. Mueller, Rajesh Menon, Erik M. Jorgensen, Jordan M. Gerton

## Abstract

Multicolor localization microscopy typically relies on sequential imaging and bandpass filters to distinguish fluorescent tags, which introduces temporal delays during live imaging, and decreases photon yield. By engineering the point-spread function (PSF), different fluors can be imaged simultaneously and distinguished by their unique patterns, without discarding photons. Here, we insert a silicon-dioxide phase plate at the Fourier plane of the detection path of a wide-field fluorescence microscope to produce distinguishable PSFs (X-PSFs) at different wavelengths. We demonstrate that the resulting PSFs can be localized spatially and spectrally using a statistics-based computational algorithm and can be utilized for hyper-spectral super-resolution microscopy of biological samples. Single PSFs in fixed U2OS cells were acquired using dSTORM with simultaneous illumination of fluors without emission filters. The modified PSF achieves ∼21 nm lateral localization precision (FWHM), ∼17 nm axial precision (FWHM) with an average of 1,800 - 3,500 photons per PSF and a background as high as 130 - 400 photons per pixel. The modified PSF can distinguish up to three fluorescent probes with ∼80 nm peak-to-peak separation between consecutive spectra.

## 1. Introduction

A driving need in cell biology is the ability to track proteins in a living sample at a scale meaningful for a protein, that is, with nanometer-scale spatial resolution and second-scale temporal resolution. In this context, Single Molecule Localization Microscopy (SMLM) [1] holds great promise and has transformed biological fluorescence microscopy over the past two decades. SMLM techniques [2] like PALM [3–6], STORM [7–9], and PAINT [10,11] have overcome the optical diffraction limit imposed by traditional light microscopes and have produced sharper images and videos of intracellular processes [12].

One challenge that arises when SMLM techniques are applied to imaging live cells with nm-scale spatial resolution is the inherent tension between obtaining high imaging speeds, which could reveal rapid structural rearrangements, and spectral resolution by efficiently separating the signal from multiple fluorophore species (fluors), which could reveal intermolecular interactions. Initially, SMLM techniques were applied to single fluor species [3,6,8], and multi-color imaging was accomplished by separating different excitation or emission spectra either in time (serial excitation/collection) [13], which slows acquisition speeds, or in space (dedicated fields of view on a camera and a system of spectral filters) [14–16], which results in the loss of signal.

One approach to improve the utility of SMLM techniques is to modify or engineer the point-spread function of the collected fluorescence emission, which has enabled, for example, increases in axial localization precision [16]–[22] and spectral identification of flour emission [16,24–31]. The PSFs are engineered by introducing an additional optic such as a grating, phase ramp, or phase mask (created by a liquid crystal spatial light modulator or dielectric) at the Fourier plane of the detection path [4,19,32]. Statistical approaches such as maximum likelihood estimation [33] and deep learning algorithms [34–37] have been used to localize the engineered PSFs with a precision of tens of nanometers. PSF engineering has yielded both wavelength and position-dependent PSFs [38,39] and may thus eliminate the need to use emission filters to distinguish between different fluor species. This could improve SMLM by reducing the photon loss incurred by the filters, which can increase image acquisition speed and spatial resolution. Similarly, wavelength-dependent PSFs may also obviate the need to use conventional spectrometers for spectral identification, which are fairly incompatible with wide-field single-molecule imaging techniques such as SMLM. In addition, engineered PSFs have the potential to distinguish between highly overlapping spectra that cannot otherwise be separated using emission filters. For example, a single laser could be used to excite different proteins tagged with only red dyes which could be distinguished by their unique PSFs. Spectral unmixing [40], biplane imaging [5], and spectral multiplexing [41] are compatible and complementary alternatives to PSF engineering.

In this paper, we modify the PSF in a spectrum-dependent manner using a phase mask created by a customized glass phase plate (Fig. 1) and demonstrate nm-scale spatial localization and simultaneous differentiation of up to three fluor species. The modified PSF is referred to as an “X-PSF” based on its distinctive shape and the customized phase plate the “X-PP” as explained below. By inserting the X-PP into the Fourier plane of a commercial fluorescence microscope, we demonstrate spectral differentiation under simultaneous illumination without using spectral filters, and successfully image and identify microtubules, mitochondria and ds-DNA in a fixed cell. Compared to some of the other PSF-engineering approaches mentioned in the introduction, the X-PP is easy to manufacture at scale using standard masking and chemical vapor deposition technology. Furthermore, the X-PSF is compact since the X-PP has only four quadrants (Fig. 1), which can ultimately lead to larger signal-to-noise ratios per pixel since it is not spread out over as many pixels on the CCD camera detector. Additionally, since the difference in thickness of the XPP quadrants is less than 1 micron, it does not scatter much of the incident light, leading to relatively little photon loss. Other advantages come from the simplicity and transparency of using a maximum likelihood estimation fitting algorithm rather than an AI-based deep learning approach. The maximum likelihood approach is based on a physical model with realistic and easy-to-interpret parameters, and it enables one-shot fitting of the position (in 3D) and wavelength of each localized fluor. One disadvantage of the X-PSF is that the imaging depth along the optic axis is limited to a range of ∼1 micron due to crosstalk between the axial position and spectral fitting parameters.

**Fig. 1.**
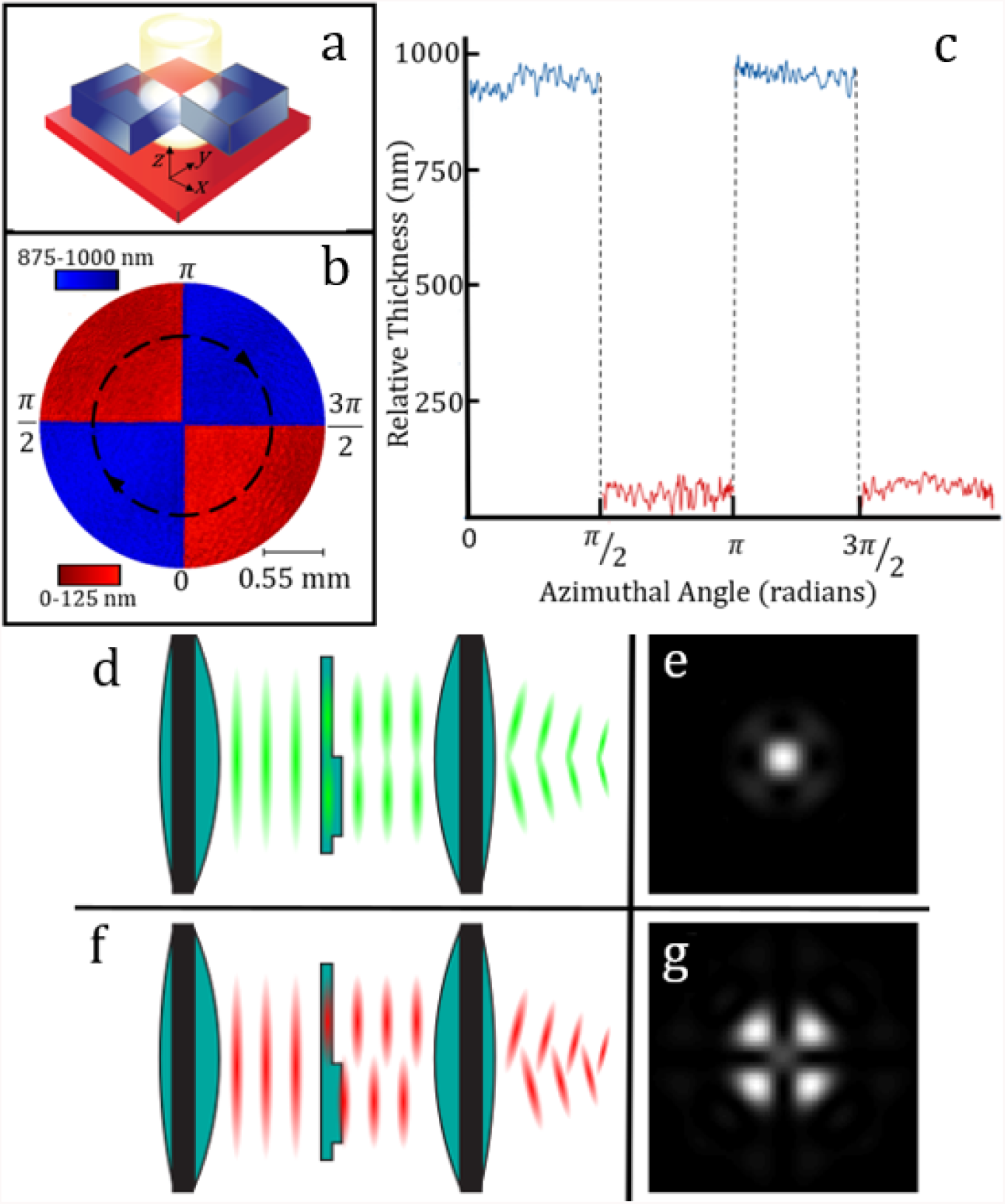
Geometry and working principle of the phase plate. a) A schematic of the phase plate. The blue regions are SiO _2_ depositions on top of a thin glass plate colored red. The emission light beam is incident on the phase plate at its center as indicated by the bright spot. b) A color-coded height map of an approximately 2 mm diameter section of the phase plate’s center in the *x-y* plane measured by a Twyman-Green interferometer, as a thickness difference from a nominal zero. c) The height profile (z) along the dashed black circle. The working principle of our phase plate is shown in panels d) to g). Panels d) and f) show how the phase plate affects fluorescence light of different wavelengths (500 and 680 nm respectively). e) and g) show the respective simulated PSFs.

## 2. Methods

### 2.1 Working Principle

The phase plate’s geometry consists of four quadrants, with each diagonal pair having the same thickness (Fig. 1(a)). We measured the surface profile of the phase plate using phase-shifting interferometry with a Twyman-Green setup [42,43] as shown in Fig. 1(b-c). The quadrants indicated in blue are Δt = 960 ± 20 nm thicker than those in red, which leads to variations in the optical path length, producing wavelength-dependent PSF profiles as illustrated in Fig. 1(d)-(g). Fig. 1(d) and (f) depict collimated light propagating through the phase plate, where the top half of the phase plate is drawn as a thin quadrant. As collimated fluorescence passes through the thin quadrant, it acquires a smaller phase shift than the light passing through the thick quadrant. For green light (λ = 500 nm), the phase shift is about 2π, leaving the wavefront effectively flat, and producing a PSF at the detection plane that is similar to an Airy disk, as shown by the simulation in Fig. 1(e). For red light (λ = 680 nm), a phase shift of about π occurs (Fig. 1(f)), resulting in destructive interference at the center of the PSF, which produces the cross-shaped null shown in panel (g). Hereafter, we refer to the PSF shapes in panels (e) and (g) as the canonical forms of our PSF. For wavelengths between canonical values, the PSF is a superposition of the two canonical forms in different proportions depending on the wavelength. As the wavelength increases from green to red, the intensity migrates outward along crossing diagonal lines from the center of the PSF. Thus, we refer to this PSF as the “X-PSF”, and the phase plate as the “X-phase plate” (X-PP).

### 2.2 Optical Setup

The X-PP was manufactured using a plasma-enhanced chemical vapor deposition process. More details about the design and fabrication of the X-PP and its mount can be found in the supplementary material (see Supplement 1). Fig. 2(a) shows a greatly simplified diagram of the detection arm of our microscope. Two parallel bundles of rays are depicted, which pass through the X-PP from two different emitters on the sample. The parallel bundles of rays overlap at the Fourier plane. It is important to place the X-PP at this location, as this renders the X-PSFs translationally invariant with respect to both the sample and image (camera) planes [22], [40]. The phase plate was manually adjusted to be as close as possible to the Fourier plane (given constraints of the microscope’s optical path, we estimate within ∼5 mm), and to align the center of the X-PP (i.e., where the quadrants meet) with the optic axis until the X-PSFs appeared symmetric on the camera. X-PP alignment was quick, taking only about 15 minutes before each imaging experiment. X-PSFs preserve their pattern when the X-PP is rotated around the optic axis. This property simplifies X-PP alignment and makes the differentiation between X-PSF patterns generated by different probes fairly robust against misalignment.

We conducted experiments on two different microscopes: The first microscope was a home-built wide-field inverted fluorescence microscope with four laser lines at 405, 488, 561, and 647 nm. These lasers were co-aligned using successive long-pass dichroic filters passing through an acousto-optic tunable filter (AOTF) to control the intensity of each. After the AOTF, all laser lines were coupled into a single multimode patch cable to transfer the laser power from our laser bench to the bench on which the microscope was constructed. A detailed diagram of this setup is given in the supplementary material (see Supplemental 1). However, it is functionally similar to the diagram in Fig. 2, representing the layout of the second microscope – a Vutara 352 commercial microscope. For a proof of principle experiment using fluorescent beads, a water-immersion objective with a numerical aperture of 1.2 was used, whereas for imaging biological samples, a silicon-immersion objective with a numerical aperture of 1.3 was used. In both microscopes, we used the X-PP at a Fourier plane in the collection path to modify the PSF to be sensitive to different spectra.

**Fig. 2.**
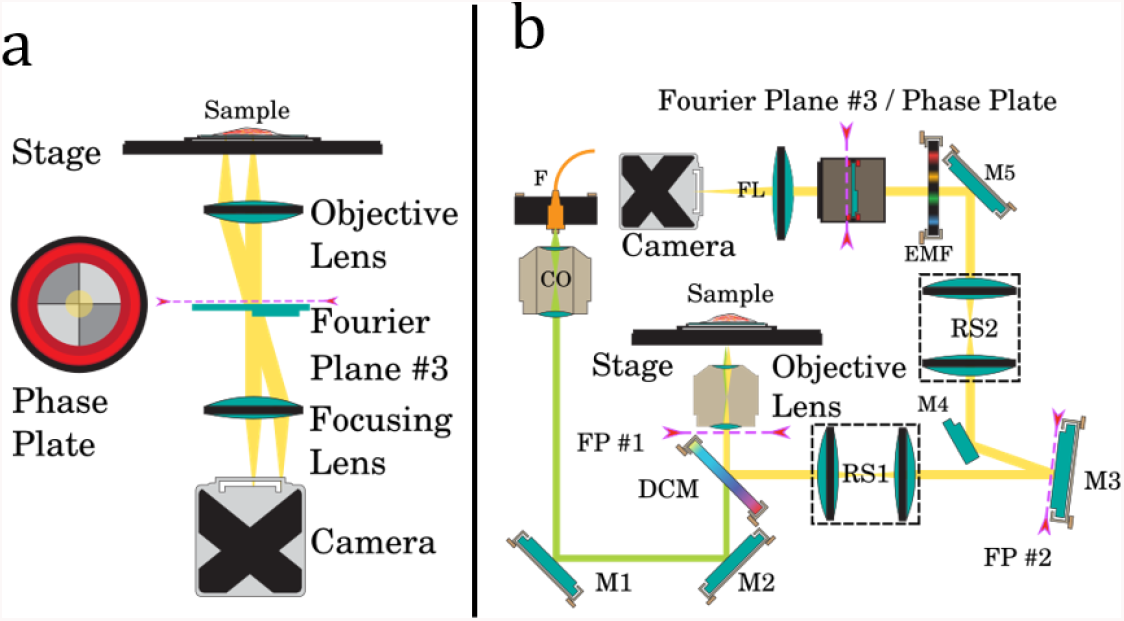
Optical setup of the microscope. a) A very simplified diagram of the detection path of our microscope. b) A more detailed diagram of a Vutara 352 super-resolution microscope. Fourier planes are indicated by magenta dashed lines. Labels are as follows: F: fiber out-couple; CO: collimation objective; M1, M2, M3, M4, M5: silver mirrors; DCM: dichroic mirror; FP #1, FP #2 Fourier planes; RS1, RS2: relay lens systems; EMF emission filter; FL focusing lens.

### 2.3 Numerical Analysis

We wrote a custom library of classes and scripts in MATLAB to analyze our raw multi-color localization microscopy X-PSF data as described below. Here, we localized the PSFs in three spatial dimensions and sorted them by the fluor type using a maximum likelihood estimation approach by comparing experimental X-PSFs with X-PSFs constructed via a mathematical model.

#### 2.3.1 PSF Model

Although many others have modeled the PSF by vector-based diffraction [26,27], scalar diffraction theory was sufficient to describe the X-PSF. We modified the Gibson-Lanni PSF model as described by Kirshner [41], and Born and Wolf [42] with a term representing the phase pattern imparted by the X-PP to fit the signal PSFs in our data:

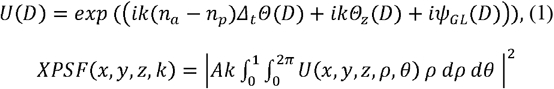

where Ψ_GL_ represents the standard phase terms in the Gibson-Lanni PSF model; Δ_t_ is the difference in thickness between the thick and thin regions of the phase plate; n_a_ and n_p_ are the indices of refraction for air and the phase plate, respectively; Θ describes the topography of the phase plate, and its maximum value is normalized to unity; and Θ_z_ describes the optical aberrations incorporated into the X-PSFs from the microscope modeled as weighted Zernike polynomials. The integral is taken over the unit disk, D, which represents the illuminated portion of the Fourier plane. A is a normalization constant such that the integral of the X-PSF is unity. Finally, (*x, y, z*) is a point in the image plane of the microscope and *k* is the wavenumber. The parallel bundles of rays from emitters away from the focal plane make very small angles with the phase plate hence the term identifying the phase contribution from the X-PP is still a good approximation for off-plane emitters as well. The X-PSF model is elaborated in the supplemental material (see Supplemental 1).

Fast-Fourier transforms (FFTs) can efficiently calculate a simplification of the diffraction integral representing light propagation of a PSF-engineered microscope [47]. However, we found that padding the numerical representation of the Fourier plane with zeros to increase the FFT sample rate to match the camera’s pixel size was too slow for our analysis goals. Instead, we calculated the diffraction integral using a Gaussian cubature formula of the unit disk as given by Cools et al. [48] with the topography of the phase plate being modeled using sigmoid functions.

Accounting for the optical aberrations inherent in our microscope was essential for matching the measured X-PSF and our model X-PSF. We account for the optical aberrations by incorporating a weighted sum of Zernike polynomials (Θ_z_) - horizontal and vertical tilt, oblique astigmatism, vertical and horizontal coma, and primary spherical aberration - in the diffraction integral (Eq.1). Experimentally, we ensure the microscope is properly focused using 150-200 nm diameter fluorescent beads, so we didn’t include defocus terms in the Zernike polynomials; furthermore, we employ single-plane imaging, so we left out multiplane terms from Kirshner et al. [45]. Before imaging biological samples, we extracted the Zernike weights characteristic of the microscope by acquiring multiple X-PSFs from fluorescent beads (FluoSpheres Tetraspeck beads from ThermoFisher) illuminated with 488, 561, and 647 (or 640) nm laser lines (Yellow-Green, Orange, and Dark-Red emission spectra). In this calibration, we acquired a stack of images at different depths (*z*) within the bead sample and used a maximum likelihood estimation algorithm (see below) to fit the model X-PSF to these experimental X-PSFs. For the fitting, we used a monochromatic representation of the X-PSF corresponding to the peak wavelength of the emission spectra after smoothing the model X-PSF image by Gaussian blurring. Regions of interest (ROI) were selected using a Python blob detection algorithm (Difference of Gaussian method) [49]. The X-PSF intensity distribution is at most 30% broader than the standard Airy PSF for emitters close to the focal plane, which is equivalent to about one camera pixel around the periphery of the PSF; for emitters far away from the focal plane, the breadth of the intensity distribution of the X-PSF is almost the same as for an Airy pattern (see Supplemental 1).

#### 2.3.2 Maximum Likelihood Estimation

With a model of the X-PSF in hand, emitter localization and wavelength estimation are done in two steps. The first step is to guess the emitter wavelength, lateral position on the camera, and depth (axial position *z*) of each experimental X-PSF in the field of view. We make this guess by constructing a lookup table using the Zernike-calibrated X-PSF model. The lookup table has four dimensions: *x, y*, and *z* spatial coordinates and the spectral coordinate λ. The spatial dimensions of the lookup table consist of the X-PSF model at positions separated by 50 nm in-plane and axially, while the spectral dimension consists of two (or three) monochromatic representations of the calibrated X-PSF.

The initial guess becomes the starting point for the second localization step in which the negative log-likelihood between the data and the model is minimized using a Nelder-Mead simplex algorithm [50]. This second step is done with all four dimensions of the X-PSF (*x, y, z*, and λ). During this part of the analysis, a monochromatic model of the X-PSF is again used because fitting using the Nelder-Mead algorithm with a polychromatic representation of the PSF is too computationally costly. A Gaussian smoothing filter is applied to the Zernike-calibrated monochromatic X-PSFs modeled at the emission peak of each spectrum. We found the Nelder-Mead algorithm to be computationally faster than the Newton-Raphson method because the latter required calculating the Hessian of the objective function. The algorithm records a PSF’s spatial and wavelength coordinates after each fit had gone through 3,000 iterations and we bin the localizations considering the fitted wavelengths and then use a hard threshold to determine the associated probe type. The result was passed to the rendering software (Vutara SRX) to produce images with varying statistical filtering criteria. These filters were applied manually by the user to qualitatively optimize the reconstructed super-resolution image.

### 2.4 Sample Preparation and Imaging

#### 2.4.1 Fluorescent Bead Sample Preparation

The samples imaged in Fig. 3 and Fig. 4 were made by diluting ThermoFisher FluoSphere beads (see results for more specific information) in a 1:10 ^4^ ratio from a stock solution in ultra-pure water. A microscope slide was prepared with 15 µl of poly-L-lysine as an adhesive for the beads and was allowed to stand on the slide for ten minutes before the excess was pipetted off. 20 µl of the dilute bead solution was then pipetted onto the slide and dried in a vacuum chamber. Once dry, the area containing the beads was secured with a coverslip using nail polish as a sealer. The sample was mounted on the microscope such that the laser light is incident through the coverslip (#1.5H).

**Fig. 3.**
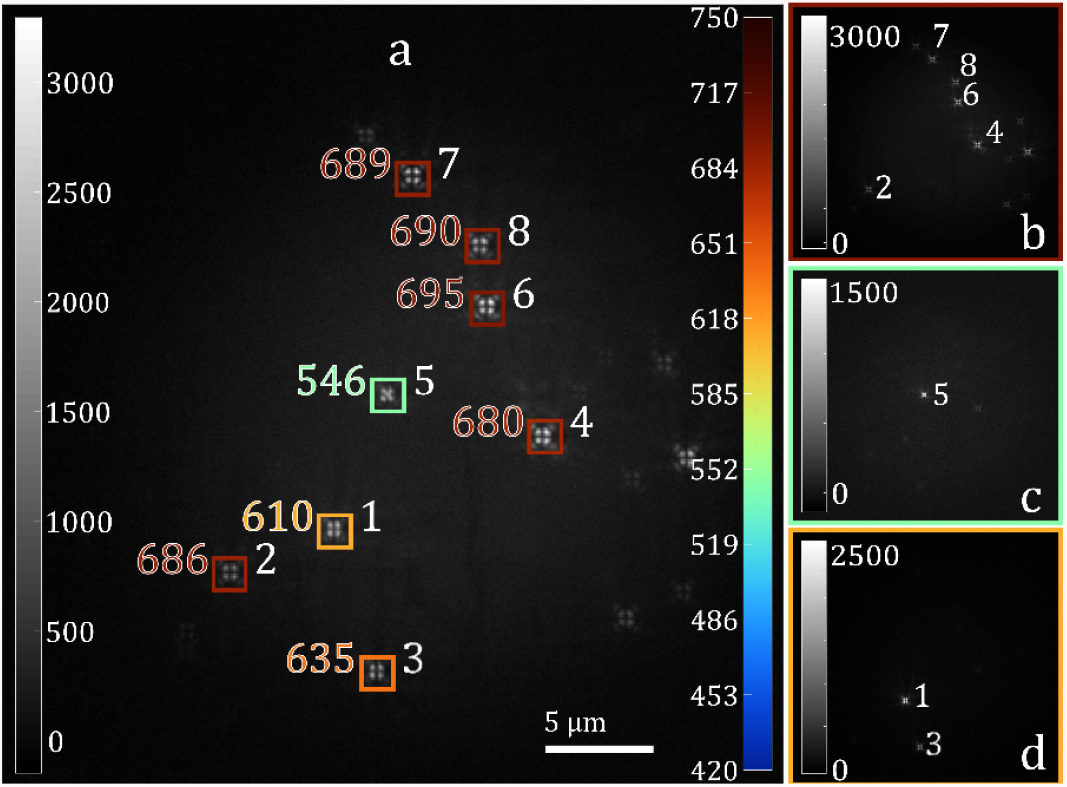
Demonstrating the ability to distinguish different spectral species of fluorescent beads using only the shape of the X-PSF. a) Three different species of beads imaged simultaneously. Colored boxes are drawn around each PSF that was subjected to the localization with a number indicating the wavelength to which each was localized. The left gray-scale bar indicates photon counts, and the right color bar indicates wavelength. Each PSF is numbered 1 − 8. b)-d) The same sample illuminated only with 647, 561, 488 nm respectively.

**Fig. 4.**
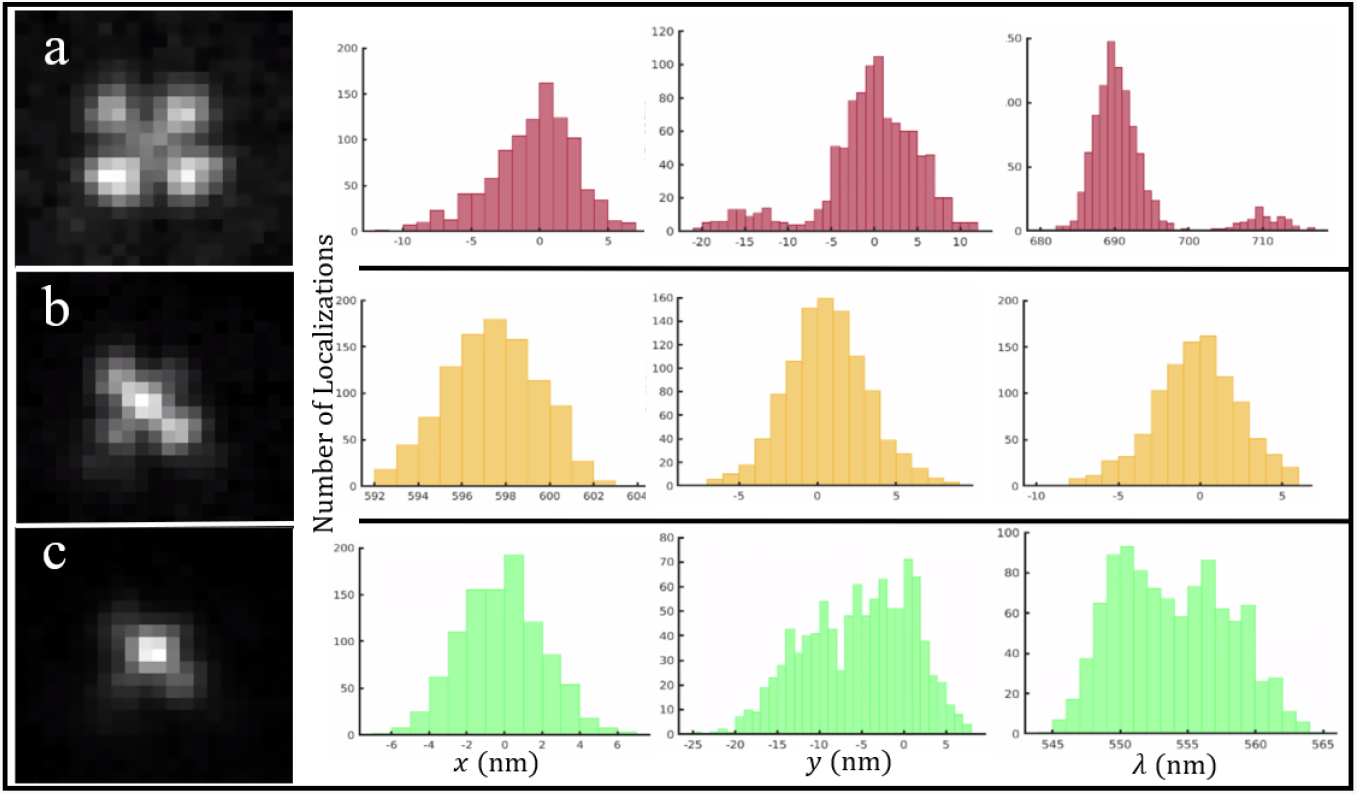
Simultaneous spatial and spectral localization of three different fluorescent spectra. a) The X-PSF of a TetraSpeck bead illuminated by the 640 nm laser line and its binned localizations for 1,000 camera frames. b) and c) The PSFs of the same bead illuminated by 561 and 488 nm laser lines separately and their corresponding histograms similar to (a). The *y* and λ localizations show two peaks for the 641 nm laser line (Dark-Red) but one peak is only 10% of the other and the ratio between the histograms areas too is 1:10.

#### 2.4.2 Biological Sample Preparation

U2OS cells grown in #1.5H Ibidi chambers (µ-Slide 8 well Cat#80826) at 37°C in 5% CO_2_ airgas were fixed in 37°C 3% paraformaldehyde (EMS) and 0.1% glutaraldehyde (EMS) in PEM buffer (100 mM PIPES, 1 mM EGTA, 1 mM MgCl_2_, pH 7.0) for 15 minutes. Glutaraldehyde autofluorescence was quenched by adding fresh 0.1% sodium borohydride in PBS for 7 minutes. Cells were blocked and permeabilized in blocking buffer (5% donkey serum, 0.1% Saponin (w/v), 0.05% Triton X-100 in PBS) for 45 minutes at room temperature. Cells were incubated within blocking buffer rabbit anti-Tomm20 (abcam AB78547) and mouse anti-HSP60 (R&D systems MAB1800) (Thermo-Fisher) or with mouse anti-alpha tubulin (Sigma) and rabbit anti-detyrosinated tubulin (Sigma). Cells were incubated overnight at 4°C with shaking. Cells were washed 3 to 5 times in PBS. Cells were incubated with secondary antibodies anti-rabbit Alexa Fluor 647 (Thermo-Fisher), anti-mouse CF568 (Biotium), and Phalloidin-Alexa Fluor 488 (Thermo-Fisher) for 1 hour at room temperature. Cells were washed 3 to 5 times in PBS. Cells were postfixed in 4% PFA for 5 minutes at room temperature and washed 3 to 5 times in PBS and cells were imaged in standard Gloxy imaging buffer for dSTORM: 20 mM cysteamine, 1% (v/v) 2-mercaptoethanol, and oxygen scavengers (168 AU glucose oxidase and 1404 AU catalase) in 50 mM Tris, 10 mM NaCl buffer with 10% w/v glucose at pH 8.0.

### 3 Results

### 3.1 Ability to Distinguish Between Spectra

To validate our approach to spectrally sensitive PSF engineering, we used a phase plate with a 660 nm step height to verify the ability of the X-PSF to distinguish between spectra; this X-PP (X-PP1) was similar to the one shown in Fig. 1 and generates PSFs in the X-PSF family. We illuminated three different species of fluorescent beads (ThermoFisher FluoSpheres: Dark-Red, Orange, and Yellow-Green, prepared as described above) with 647, 561, and 488 nm laser line sequentially at first to identify individual beads by type and then simultaneously to collect experimental X-PSFs for our MATLAB localization algorithm. For both illumination schemes, we collected ∼6,000 photons for each X-PSF. By comparing the wavelengths estimated using our MATLAB algorithm, indicated by the color-coded wavelength values and boxes in Fig. 3(a) (simultaneous illumination) to the control panels (b)-(d) (single line illumination), it is clear that the estimation program correctly identifies the different fluorescent bead species near the axial focus.

### 3.2 Localization Precision

We quantified the simultaneous spectral and spatial localization capability of the X-PSFs using our localization algorithm by repeating the same procedure described above for X-PSFs with three different bead colors, as shown in Fig.4(a) for 1,000 camera frames. For this experiment, we used X-PP2 with a step height of 960 nm. The histograms for localizations in *x, y*, and λ for the three bead spectra are shown in Fig.4 (b)-(d). Compared to X-PP1, X-PP2 was more sensitive to changes in the wavelength and the axial (*z)* spatial coordinate (i.e., the PSF pattern was easier to differentiate) and therefore was used for biological imaging. The family of X-PSFs produced by X-PP2 showed a lateral spatial localization and a spectral localization precision of ≤10 nm (FWHM) near the focal plane for Dark-Red (∼16,000 photons per PSF and ∼50 background photons per pixel) and Orange (∼22,000 photons per PSF and ∼50 background photons per pixel) PSFs. For Yellow-Green PSFs (∼17,000 photons per PSF and ∼50 background photons per pixel), the localization precisions were ∼15 nm. These photon counts are significantly higher than what is expected from individual PSFs in most biological samples, and thus these FWHM measurements represent an estimated limit to the localization precision near the focal plane.

### 3.3 Multi-Spectral Super-resolution Imaging of Biological Samples

We demonstrate simultaneous two-color and three-color dSTORM [51] imaging using the X-PSFs created by the X-PP2 phase plate placed at the last Fourier plane of the Vutara 352 microscope. The spectra were localized based only on PSF shape and without any emission filters in the collection path other than the dichroic filter used to separate the excitation light. We used 10,000 camera frames at 25 ms each to render the images. Figure 5(a) shows the *x-y* plane of a super-resolution image of microtubules composed of α-tubulin tagged with Alexa Fluor 647 (green), and TOMM20 tagged with CF568 on the outer membrane of mitochondria (red) in fixed U2OS cells. The cell sample was illuminated simultaneously by the 640 and 561 nm laser lines. The two protein species are uniquely identifiable. TOMM20 appears interwoven within the microtubule forest as expected. Figure 5(b) shows the wavelength distribution of the localization and Fig. 5(d) shows the same image in the *x-z* plane. Fig. 5(c) shows the single probe images of the segment of Fig. 5(a) that is inside the dashed rectangle. For the two-color image, we observed ∼3,500 photons per PSF on average over an average background of ∼400 photons per pixel. Fig. 6(a) shows the *x-y* plane of a three-color super-resolution image of α-tubulin tagged with Alexa Fluor 488 (blue), TOMM20 tagged with CF568 (yellow), and ds-DNA inside mitochondria tagged with Alexa Fluor 647 (red) in fixed U2OS cells. The cell sample was illuminated simultaneously by the 488, 561 and 640 nm laser lines. The ds-DNA localizations are contained inside mitochondria as expected. We observed ∼1,200 photons per PSF on average for the localizations in the green channel and ∼ 2,500 for the other two channels at a background of ∼130 photons per pixel. There is considerable cross talk between the green and the orange channels when compared to the orange and red channels.

**Fig. 5.**
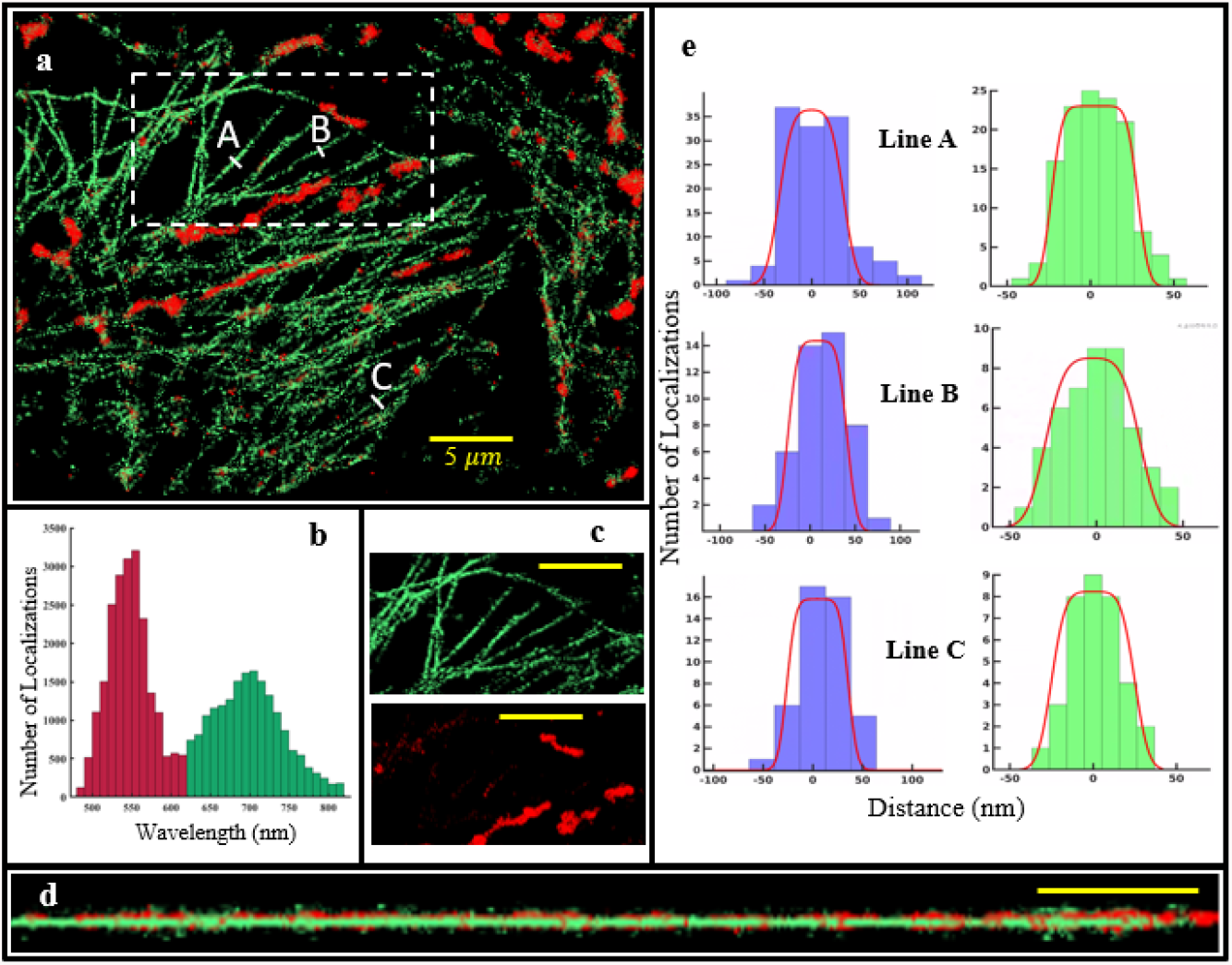
Two-color fixed cell imaging. a) Tubulin, labeled with Alexa Fluor 647, is rendered in green; TOMM20, labeled with CF568, is rendered in red. The cell sample was illuminated simultaneously by the 640 and 561 nm laser lines, and the spectra were localized based only on PSF shape without using any emission filters. The excitation intensity for the two laser lines was ∼5 kW/cm ^2^. A 405 nm laser was used at ∼1 W/cm ^2^ to control the fluorophore blinking rate by driving fluors in dark states back to the fluorescent ground state. b) The wavelength distribution of the localizations; the colors indicate the wavelength threshold for rendering in green or red. The threshold value used to separate the probes is 620 nm. c) Single probe image renditions of the rectangular segment indicated in dashed white lines in (a). d) The *y-z* cross section of a portion of (a). e) The density of localizations along A-C line segments in (a). Blue histograms indicate the density of localizations across the width (*x-y* plane) and green histograms the depth (*z*) of a microtubule respectively. The convolution of a Gaussian and a rectangular function are fitted to each histogram as shown in red. For the width (blue) histograms, the Gaussian FWHM are 25, 20 and 16 nm (A-C) and the width of the rectangular function is 68, 66 and 61 nm. For the depth (green) histograms, the Gaussian FWHM are 12, 20 and 15 nm A-C and the width of the rectangular function is 52, 55, and 50 nm. All the scale bars in (a)-(c) represent 5 μm.

**Fig. 6.**
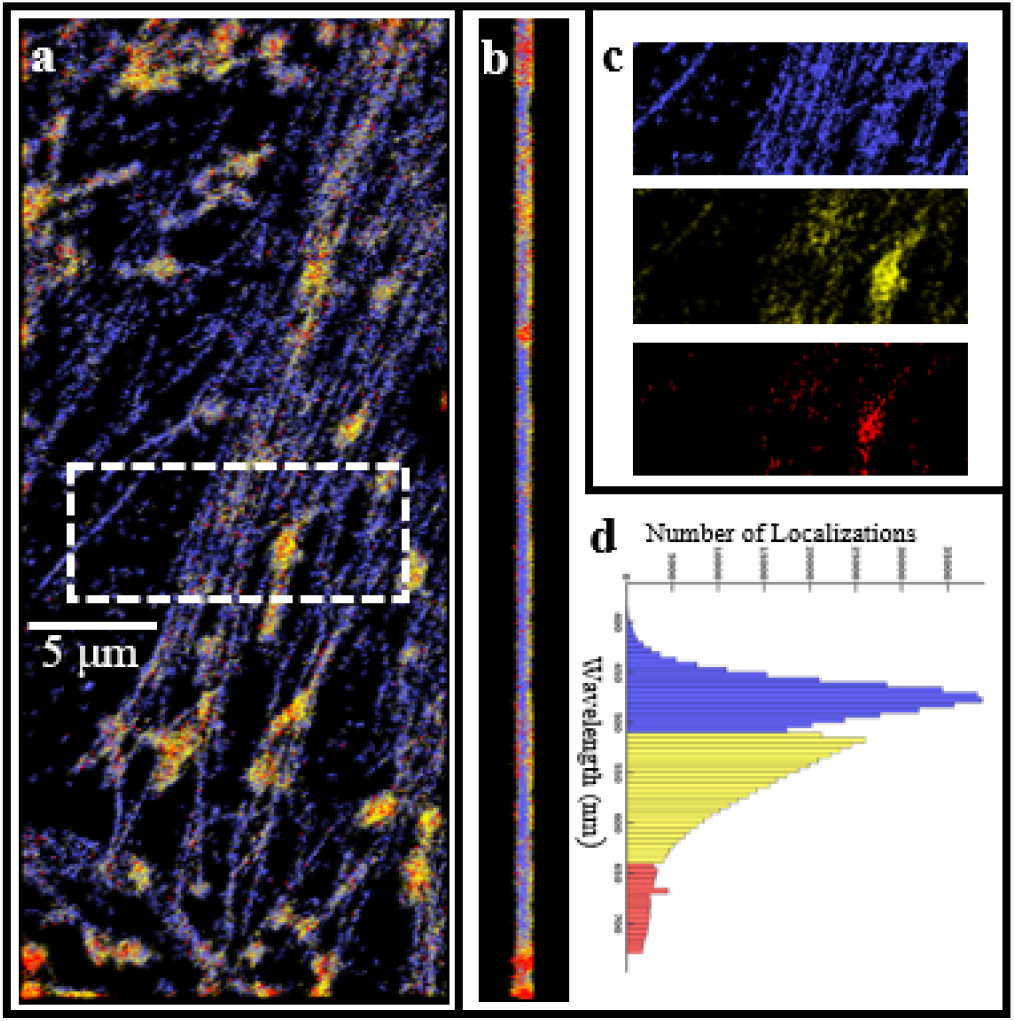
Three-color fixed cell imaging. a) Tubulin, labeled with Alexa Fluor 488, is rendered in blue; TOMM20, labeled with CF568, is rendered in yellow; ds-DNA, labeled with Alexa Fluor 647 is rendered in red. The cell sample was illuminated simultaneously by the 488, 561 and 640 nm laser lines, and the spectra were localized based only on PSF shape without using any emission filters. The excitation intensity for the three laser lines was ∼3 kW/cm ^2^. A 405 nm laser was used at ∼1 W/cm ^2^ to control the fluorophore blinking rate by driving fluors in dark states back to the fluorescent ground state. b) The *y-z* cross section of a portion of (a). c) Single probe image renditions of the rectangular segment indicated in dashed white lines in (a). d) The wavelength distribution of the localizations; the colors indicate the wavelength threshold for rendering in blue, yellow or red. The threshold values separating the probes are 510 and 640 nm. All the images in (a)-(c) have a common scale bar of 5 μm included in (a).

Microtubules provide an excellent reference to test the localization precision because of their extended linear nature and their known width. Native microtubules are 25 nm in diameter [52], and antibody-coated microtubules have been measured at 60 nm by electron microscopy [53]. The histograms Fig. 5(e) show the density of localizations across three microtubule strands labeled A, B, and C in Fig.5(a). We fitted a convolution of a Gaussian and a rectangular function (indicating the finite width of an antibody-stained microtubule) to each histogram. To estimate our localization precision, we use the FWHM of the Gaussians [13,25]. The FWHM of these Gaussian fits are 16 - 25 nm in the *x-y* plane and 12 - 20 nm in the *z* direction across a microtubule. The calculated width of an antibody-stained microtubule is 65 nm on average in the *x-y* plane and 52 nm in *z*.

## 4 Discussion

A key goal in cell biology is to observe the movement of multiple proteins in a living cell at a spatial scale meaningful to a protein – specifically at the diameter of a typical protein, ∼10 nm. Here, we observe that X-PSFs created by a four-quadrant glass phase plate can be used for multi-spectral imaging with localization precision comparable to other PSF families. The discussion below evaluates the localization precision of the X-PSF, its ability to distinguish spectra, and its utility for imaging multiple probes under simultaneous excitation and detection.

### 4.1 Localization Precision

We use the Cramér-Rao lower bound (CRLB), referred to as 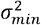 (statistical variance) in this paper, to estimate the best-case precision of the X-PSF in the *x, y, z*, and λ dimensions. We represent the center coordinates of a PSF at the image plane of the microscope as *x*_*0*_ and *y*_*0*_, the axial coordinate relative to the focal plane of the microscope (i.e., at the sample) as *z*_*p*_, and a representative wavelength for the emission spectra as λ. As others have noted [54–57], CRLB is calculated by inverting the Fisher information matrix and taking its diagonal elements as the *minimum* possible variance 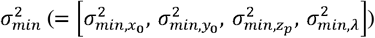 around the actual value that can be achieved when measuring the unknowns *x, y, z*, and λ. Assuming shot noise only, the Fisher information matrix I_k_ is given by Eq. 2 where 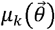is the theoretical (model) X-PSF given by Eq. 1, 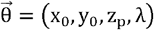are the parameters being estimated:

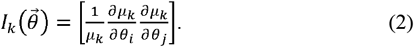

The theoretical precision in 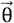for the X-PSFs calculated as the 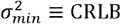at three different emission wavelengths (λ = 680, 585, and 515 nm) is shown in Fig. **7**(a)-(c) respectively. For this calculation, we neglected Zernike aberrations and assumed that light detection follows Poissonian statistics and that the background is uniform across all pixels. We observed 1,000-5,000 photons per PSF for the cells, and therefore the precisions are plotted at 2,500 photons as an average. Fig. **7** suggests that the axial localization precision is expected to be worst at the focal plane, since the X-PSFs are symmetric at the focal plane and the asymmetry that allows to distinguish between the negative and the positive axial values is small at the vicinity of the focal plane. Since the X-PSF is asymmetric far away from the focal plane, the axial localizability improves (see Supplemental 1).

### 4.2 Simultaneous Multicolor Imaging

One promise of simultaneous multicolor imaging is that it opens the possibility of imaging live cells on fast time scales. Currently, imaging of protein localizations using filter sets can require minute-long acquisitions [1–3]. Proteins and organelles can move significantly during this acquisition time. However, simultaneous imaging could potentially identify proteins as they interact or co-migrate in a cell despite the challenges discussed above. In addition, it might be possible to design a labeling scheme whereby different fluorophore species with tightly spaced emission spectra could be excited with a single laser line and still be separated spectrally using engineered PSFs. Imaging with a single laser will be less expensive and less damaging to the biological samples than illuminating with multiple laser lines (as was done here).

During simultaneous multicolor imaging using X-PSFs, all fluorophore species are illuminated simultaneously and the emission is not separated spectrally using emission filters or otherwise. In this case, the background signal can be elevated due to overlapping PSFs from the different fluor species, from the same species, or cell autofluorescence. Any of these background sources, which are more prominent in simultaneous multicolor imaging relative to single-color imaging, obscure the structure of the X-PSF and it is somewhat more difficult to identify individual emitters by their spectra. Thus, a fluorophore species with significantly weaker emission compared to others in the vicinity may get lost in the background, and it may be difficult for the fitting algorithm to identify the regions of interest and also to converge on a solution. For instance, Alexa Fluor 488 PSFs in our data acquisitions yielded only half the photons as the other two fluors. This may have contributed to the cross-talk between green and orange channels. Theoretically, limiting blinking rates and editing out poor wavelength localizations could limit such miscalls but would also reduce localization density.

Another challenge in simultaneous multicolor imaging is controlling the independent blinking rates of each fluor species. To restore fluors to the ground state, we use a 405 nm laser which illuminates all fluors simultaneously. But the photophysics of each fluor species and the label density for each probe may require a different 405 nm laser intensity to optimize blinking. In addition, highly concentrated clusters of a protein and unstable triplet states can lead to regions that require a different 405 nm laser intensity to ensure that PSFs are sufficiently sparse to be localized. Matching PSF densities from different fluors might require diluting the fluorescent labels of particular proteins.

### 4.3 Differentiating Between Probe Species Using the X-PSF

The ability of the X-PSF to differentiate between fluor species based on their emission spectra was studied using sequential imaging of a U2OS cell and Monte-Carlo simulations using experimental parameters. In Fig. 8 we show simulations of correct spectral assignments for the X-PSF as a function of the number of background photons per pixel for the three dyes Alexa Fluor 647 (tagging microtubules), CF 568 (tagging mitochondria), and Alexa Fluor 488 (tagging clathrin), and the rendered image of the sequential imaging (control experiment) to compare with the simulations. The simulated sample background is primarily caused by autofluorescence and fluorescence from out-of-focus emitters, both belonging to the same or to different probe species. We simulated noisy X-PSFs by incorporating the sample background, shot noise in photon detection, and camera readout noise to the model X-PSFs at multiple spatial coordinates for a specific probe. Furthermore, we employed the same localization algorithm used for biological data analysis (Fig. 5 and Fig. 6) to determine the probe type. We used a distribution of photons similar to the photon distribution observed in the control experiment and that of Fig. 5 with an average of ∼ 3,500 photons per PSF. The number of correct identifications is the number of X-PSFs correctly sorted by the algorithm as a percentage of the total number of the input X-PSFs belonging to that probe as a function of the number of background photons. For a simulated background of 100 photons per pixel, correct identifications for Alexa Fluor 647, CF568, and Alexa Fluor 488 probes were 95%, 89%, and 94%, respectively.

**Fig. 7.**
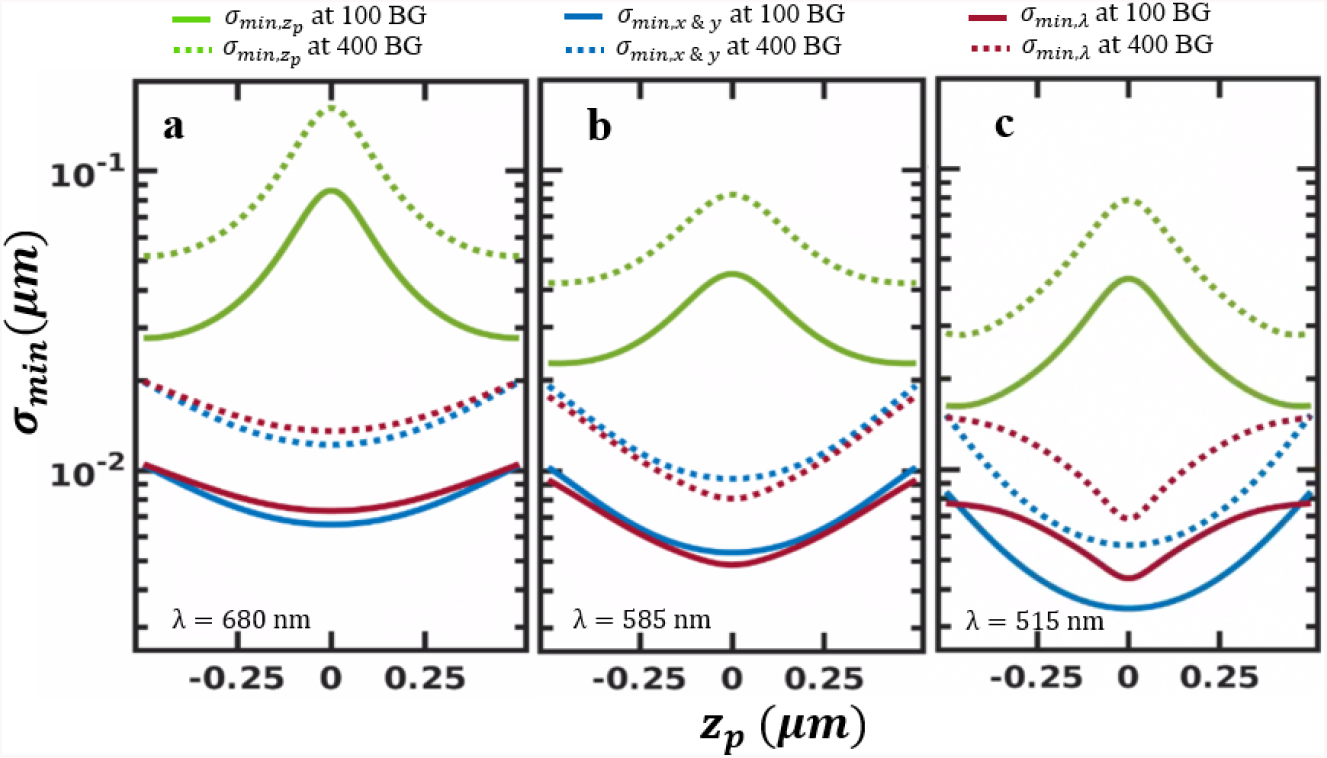
Theoretical precision of the X-PSF as a function of position relative to the focal plane, *z*_*p*_, at different wavelengths for 2,500 photons per PSF at 100 and 400 background photons per pixel. The wavelengths are 680 nm (a), 585 nm (b), and 515 nm (c). The green, blue, and red curves correspond to axial direction, the in-plane directions, and the wavelength respectively. Solid lines correspond to 100 background photons per pixel and the dotted lines to 400. The wavelengths roughly correspond to the peak wavelengths for the “Dark-Red”, “Orange”, and “Yellow-Green” FluoSpheres spectra in Fig. 4.

**Fig. 8.**
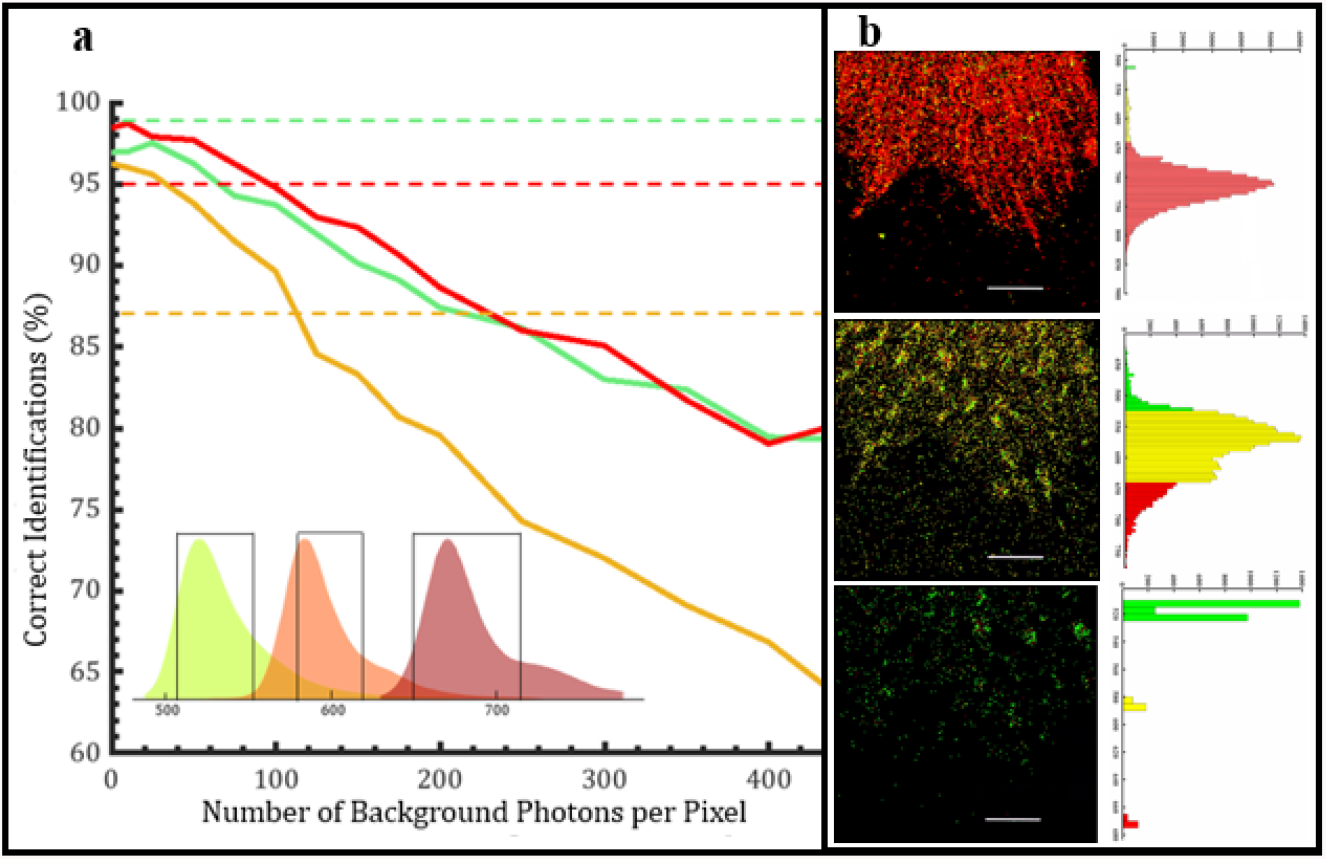
Spectral distinguishability of the X-PSFs using a Monte-Carlo simulation. a) The simulated percentage of correct spectral identifications for the probes AXF 647 (red), CF 568 (orange), and AXF 488 (green) are plotted as a function of the number of background photons per camera pixel. In the simulations, we used a distribution of photons similar to the photon distribution observed in the control experiment and the biological imaging in Fig. 5 having an average of ∼3,500 photons per PSF. The dashed lines indicate the best-case spectral discernibility using emission filters color-coded the same as the dyes. The three spectra are shown in the inset with the transmission windows of the dichroic filter denoted by the black lines. b) The *x-y* plane of the identifications for red, orange and green channels from top to bottom of the control experiment (sequential imaging of each probe and separation using emission filters). The identifications are rendered in red, yellow, and green respectively. The scale bar shows 5 μm. The corresponding histograms for the assigned wavelength are at the left of each image. The horizontal axis is the number of PSF counts and the vertical axis is the wavelength in nm. The threshold values of 525 nm and 640 nm are common to all the channels and were chosen qualitatively after observing the histograms.

We prepared fixed U2OS cells with the same three distinct fluorophore labels to reproduce the imaging conditions of biological samples; however, the labeling efficiency of this fixed sample was not sufficient to yield high-quality three-color images. We imaged this sample sequentially (not simultaneously) with three laser lines and identified the fluorophore species using emission filters. We then applied our X-PSF analysis algorithm to the same sequential data set, and extracted the fluorophore species from the shape of each PSF. For a background of ∼100 photons per pixel, identifications were correctly reported for 95%, 80%, and 89% of the localizations for Alexa Fluor 647, CF 568, and Alexa Fluor 488, respectively. Truth tables for the simulations and the control experiment can be found in Supplemental 1. Mismatches between the simulation and the actual data could be due to several factors. For example, it is difficult to precisely know the number of photons per PSF in the experimental data as has been reported elsewhere [58]. In addition, variations in the background over the region of interest and having multiple (overlapping) PSFs in the same region of interest, both of which become more prominent with simultaneous multicolor imaging, increase the probability of mismatches compared to single-color or sequential multicolor imaging.

We also compare the results of the Monte-Carlo simulations to a best-case scenario using spectral filters. In particular, we calculated the expected crosstalk between different spectral channels when using the combination of dichroic and long-pass filters in our commercial microscope (indicated by the transmission profiles in the inset to Fig. 8), and then subtracted this from unity to deduce the probability for a correct identification of fluorophore species. We did not take into account spectral or shot noise when making this best-case calculation, and we assumed that all three fluorophore species emit with the same peak intensity. The results of this calculation are indicated by the horizontal dashed lines in Fig. 8.

Nevertheless, using an algorithmic approach to localize engineered PSFs can be simpler and easier to implement when compared to deep learning methods. Unlike a LCSLM, our phase mask is a transmissive, polarization insensitive, and an inexpensive glass optic that can be easily inserted into a commercial microscope, as was done here. The X-PSF models are evaluated at only 88 points (Gaussian Cubature model) which is computationally inexpensive. The PSF shapes are mathematically calculated for each *x, y, z* coordinate combination with the prior knowledge of the emission spectra, which eliminates the need of acquiring training data for every new set of conditions. It is important to note that properly calibrating the PSF model to closely resemble the data is critical and can be somewhat challenging.

## 5 Conclusion and Future Work

A four-quadrant glass phase plate can adequately alter the structure of PSFs to differentiate three fluorescent emitters for localization microscopy. We use the scalar Gibson-Lanni model with Gaussian cubature representation of the phase plate topography and the log-likelihood estimation to localize the emitters simultaneously in *x,y,z*, and λ. Overall, our approach is adaptable to various microscopy methods and the implementation is straightforward. The X-PSF, which results from placing the phase plate in the Fourier plane of the detection arm of a microscope, allows single molecules to be localized with a precision of 21 nm laterally and 17 nm axially. The proof-of-principle measurements using dye-doped beads suggests that with sufficient signal, it could be possible to resolve 10 nm spectral differences at the focal plane using our X-PSF approach. We have demonstrated simultaneous imaging of two-color and three-color fixed U2OS cells with the X-PSF. X-PSFs may improve the capabilities of new super-resolution imaging techniques such as spectral multiplexing, and biplane imaging [40,41]. The utility of X-PSFs for simultaneous live-cell imaging may be improved further by using quantum dots that have narrow emission spectra and a large fluorescent yield [59,60]. The goal is to identify spectra at nanometer-scale spatial resolution at millisecond-scale temporal resolution, for long timespans of cellular evolution.

This work also highlights some inherent challenges associated with simultaneous multicolor imaging. In particular, the rate of overlapping PSFs increases when multiple fluor species are imaged simultaneously onto the detector; this causes an increase in the background signal for any particular PSF, and also leads to non-uniform background fields, both of which compromise the spatial and spectral localizability. In the future, we will study how these challenges could be addressed, for example, by cycling quickly between different illumination lasers and synchronized readout of corresponding camera frames or using dyes with comparable emission intensities and complementing blinking rates. While this will slow down image acquisition relative to simultaneous imaging, it should yield more precise localizations, which may be the more important factor for some experiments.

One goal for future work is to improve the model PSFs to better account for aberrations and spectral factors, making our approach more robust and precise. For this work, we qualitatively chose the thickness of the four-quadrant phase plate such that the red and green X-PSFs show canonical forms. We intend to introduce an optimization scheme for the thickness to yield X-PSFs that are more efficient for different systems such as the properties of the set of fluorescent tags, level of background, and the required imaging depth. The eventual goal is to localize multiple proteins in three dimensions in a living cell.

See Supplemental 1 for supporting content.

## Acknowledgments

The authors acknowledge Drs. Carl Ebeling and Wayne Davis for their insight and feedback, and The University of Utah nanofabrication facility where the phase plate was fabricated.

## Data Availability

Data underlying the results presented in this paper are not publicly available at this time but may be obtained from the authors upon reasonable request.

