## Supplemental Information for "Simultaneous Spectral Differentiation of Multiple Fluorophores in Super-resolution Imaging Using a Glass Phase Plate"

### 1. Phase Plate Fabrication

The phase plates were manufactured using plasma-enhanced chemical vapor deposition (PECVD). First, a layer AZ 9260 photoresist was spin-coated to a thickness of 10 microns on a cleaned boro-float glass wafer having a thickness of 1.1 mm. The photoresist was then photolithographically patterned for liftoff. PECVD via liftoff provides smoother surfaces than wet or reactive ion etching. Next, a layer of SiO_2_ was deposited at 100 °C to a thickness of 960 nm on top of the substrate and was later lifted off in areas where the photoresist remained, by submerging it inchinchin acetone and applying ultrasonic power, revealing the smooth glass wafer surface. The wafer was then diced into individual chips. A chip was about 1x1 cm having a margin of about 2 mm around the deposition. The process is illustrated step by step in Fig. S 1**.**

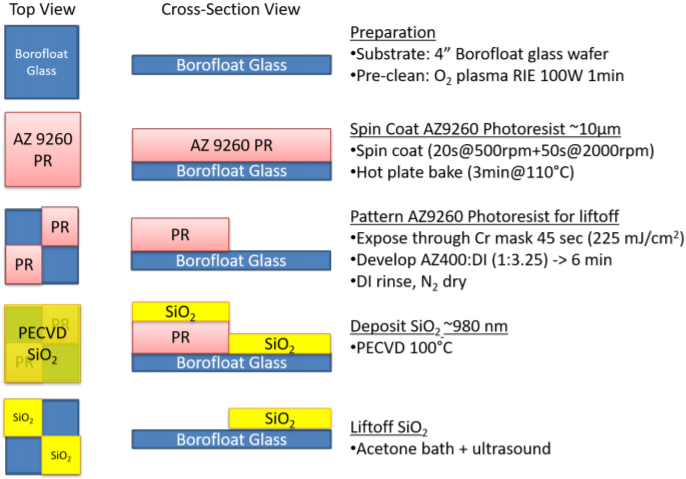

Fig. S 1 PECVD process flow. The left column shows the top view of the phase plate after each process step, the middle column shows the side view and the right column describes the process.

The phase plate mount shown in Fig. S 1 was 3D printed as an assembly of two circular caps with a smaller circular aperture at the center. Both the caps fit inside a 1-inch Thorlabs lens tube with very less wiggle space. One of the caps has a square groove at its center surrounding the aperture, to hold the phase plate at the center of the aperture. Once the phase plate sits inside the grooves of the cap, we close it with the other cap, mount it to a lens tube,module and secure it with a lens ring.

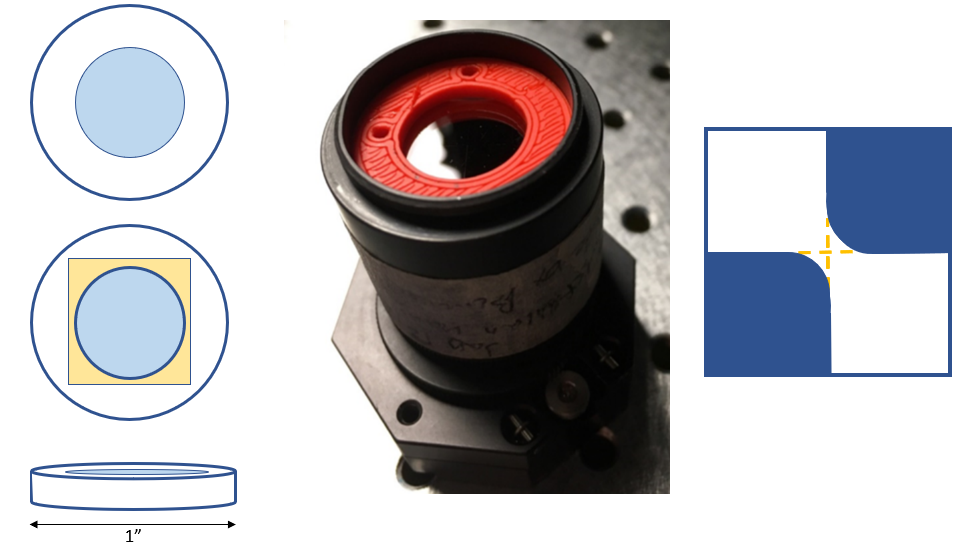

Fig. S 2 The 3D printed phase plate mount. The schematics on the left correspond to cap one and cap two when viewed from above and the side view of each cap from top to bottom. The blue regions indicate the apertures where light could pass through and the yellow region to the thin groove that secures the phase plate. The 3D printed caps are in orange in the actual image of the phase plate module in the center. A simplified drawing of the top-down view of the actual phase plate at its center is on the left. The blue area corresponds to the thick regions and the white to the nominal zero regions. The yellow dashed lines indicate the expected crossover.

### 2. Microscope Optical Setup

The home-built microscope setup shown in Fig. S 3 The home-built microscope optical setup. functions very similarly to Vutara. The optical path starts at the fiber in-couple F1 showing the large core-diameter ThorLabs patch cable which couples in three laser excitation lines: 488, 561, and 647 nm. O1 is an air objective that collimates the light introduced at F1. M1 mirror directs the excitation light to D, a quad-band pass dichroic mirror (Di01-R405/488/561/635 from Semrock). The dichroic mirror directs the excitation light through a transfer system composed of L1 and L2 converging lenses to the main objective O2. M2 and M2 are silver mirrors. The

sample S is mounted on a motorized stage, such that the detected fluorescence travels back along the excitation path, is passed through the dichroic mirror, and passes through a second transfer 4f system composed of L3 and L4 converging lenses. The signal then passes through the phase plate mounted in the phase plate module, collectively denoted by P. The phase plate rests in a Fourier plane. M4 mirror directs the light through the focusing lens L5 and a final quad band excitation filter (FF01-446/523/600/677 from Semrock), Q, that complements the dichroic mirror, onto a Photometrics EMCCD camera C. The data is then digitized and available for computational analysis.

The phase plate was mounted at the last Fourier plane of the microscope. The Fourier plane was determined by finding the physical location where the fluorescence from a dense bead sample creates the most condensed and the sharpest pattern after the L4 lens. Once the phase plate was mounted at the Fourier plane, it was moved in the perpendicular plane to light propagation using the alignment screws of the optical mount until the emission beam was at the center of the phase plate, creating a symmetric PSF shape on the camera. Lastly, the phase plate module was rotated around the light propagation axis until the four quadrants of the phase plate aligned with the *x-y* axes of the camera.

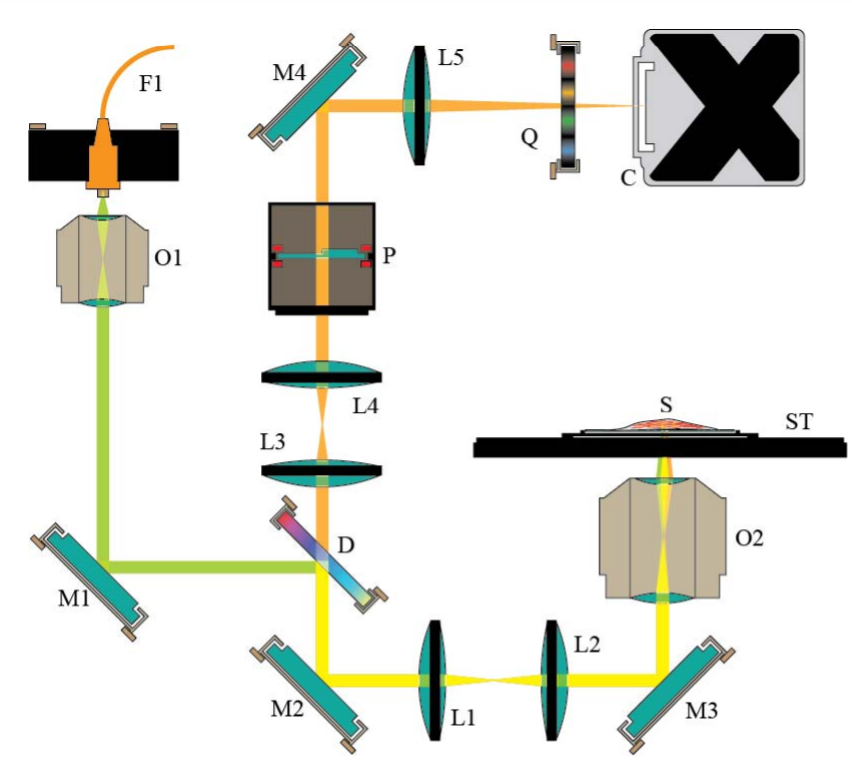

Fig. S 3 The home-built microscope optical setup. L1-L4 are lenses, M1-M3 are mirrors, D is a dichroic filter, O1, O2 are objectives, P is the phase plate module, F1 is the fiber optic cable, Q is an emission filter, C is the camera, S is the sample and ST is the sample stage. The excitation path is in yellow and the collection path is in orange.

### 3. Gibson-Lanni XPSF Model

The Gibson-Lanni XPSF model [1], [2] is described by the following set of equations. “XPSF” refers to the intensity at an arbitrary point (*x, y, z*) relative to the optical axis on the image recorded on camera for an XPSF corresponding to the wavenumber *k*. Let us define a cylindrical coordinate system by ($\rho,\theta,z$). Then, following Born-Wolf interpretation and Gibson-Lanni simplification for Kirchhoff’s diffraction integral,

$XPSF\left( x,y,z,k \right)=\left| Ak\int_{0}^{1} \int_{0}^{2\pi} U\left( x,y,z,\rho,\theta\right) \rho d\rho d\theta\right|^{2}$	Eq S 1

$$D\equiv(x,y,z,\rho,\theta)$$

$$U\left( x,y,z,\rho,\theta\right)=exp(\left( ik(n_{a}-n_{p})\Theta_{p}(D)+ik\Theta_{z}(D)+i\psi_{GL}(D) \right))$$

$\psi_{GL}\left( D \right)=k\Delta_{t}n_{i}\sqrt{1-\left( \frac{NA \rho}{n_{i}} \right)^{2}}+kz_{p}n_{s}\sqrt{1-\left( \frac{NA \rho}{n_{s}} \right)^{2}}+\frac{NA \rho}{M}\sqrt{\left( x-x_{0} \right)^{2}+\left( y-y_{0} \right)^{2}}\cos\left( \theta-\tan^{-1} \left( \frac{x-x_{0}}{y-y_{0}} \right) \right)$*.*

The terms in Eq S1 are described in Table S 1. If we were to adapt this model to a biplane or a multiplane imaging system, we need to add a defocus term as described by Kirshner et al [1] for the XPSFs corresponding to other planes, given by, $k\left( \frac{NA}{M} \right)^{2}\Delta_{z}$ , in which $\Delta_{z}$ is the axial displacement of the n^th^ plane to the focused plane.

Table S 1 The terms used in equations for the Gibson-Lanni XPSF

| **Term** | **Description** |
| --- | --- |
| $x_{0}, y_{0}$ | Lateral coordinates of an emitter in the sample domain |
| *NA* | Numerical aperture of the objective/imaging system |
| $n_{i}$ | Refractive index of the immersion medium |
| $n_{s}$ | Refractive index of the sample |
| $n_{a}$ | Refractive index of air (or else the refractive index of one of the diagonals of the phase plate) |
| $n_{p}$ | Refractive index of glass (or else the refractive index of one of the diagonals of the phase plate) |
| *M* | Magnification of the microscope |
| $\Delta_{t}$ | Sample stage displacement with respect to the working distance of the objective |
| *A* | Normalization constant such that the integral is unity |
| $\Theta_{p}$ | Thickness profile of the phase plate |
| $\Theta_{z}$ | Weighted sum of Zernike aberrations |

Since we place the phase plate at the Fourier plane of the collection path, a collimated bundle of rays would be incident on the phase plate for every emitter at the focal plane. Then, considering the geometry of the phase plate, the path difference incurred by the phase plate is $w=\frac{n_{p}\Theta_{p}}{\cos\alpha_{1}}-\frac{n_{a}\Theta_{p}}{\cos\alpha_{2}}$ where $\alpha_{1}$ and $\alpha_{2}$ are the angles that the light rays would make with the horizontal axis at the glass-air interface. However, for our imaging system, $\alpha_{1}$and $\alpha_{2}$ are very small (< 10°), and thus, $w\approx(n_{a}-n_{p})\Theta_{p}$ with a phase error of < 2%. For emitters away from the focal plane, the light rays incident on the phase plate are no longer collimated. However, for our system with light beams making small angles at the phase plate, the above approximation still produces a sufficiently accurate model for the off-plane XPSFs, facilitating *z* localizability as well.

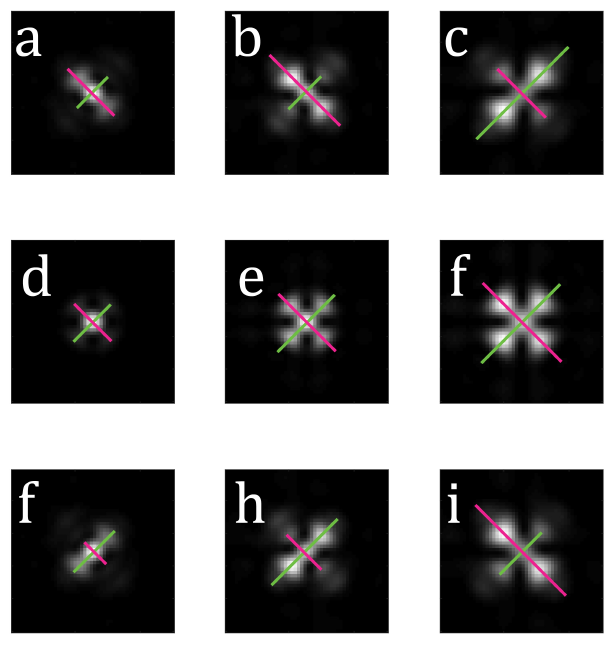

Fig. S 4 Simulated X-PSFs at different focal depths (rows) and spectra

(columns). a) XPSF for a green channel at -300 nm. As the simulated emitter

moves in focus, panel (d), and then above the focus, panel (f) (+300 nm), the PSF moves in

toward its center along the magenta line and out away from its center along the green

line. Panels (b), (e), and (h) show a similar behavior but more of the PSF is confined to the four

outer-lobes, because this PSF was calculated for the orange channel. Finally,

panels (c), (f), and (i) show the same basic behavior, however the direction of the ”tilt” left or

right of the PSF out-of-focus is opposite to that of the other two spectral channels. This is

not always the case and depends on the thickness of the X-Phase plate step, which for this

simulation was 640 nm.

The behavior of the XPSF family is illustrated using simulated PSFs in Fig. S 4 and thus explains the name “XPSF”; The intensity of the XPSFs move along the diagonals for out of focus emitters, making the shape of the letter ‘X’. For the 960 nm phase plate used for biological imaging, the direction of the “tilt” changed when moving from above and below focal plane emitters was the same for all emitters. The choice of the thickness of the phase plate may depend on the wavelengths of the emission spectra. We chose 960 nm so that the red and the green dyes we used had PSFs close to the two canonical shapes (described in the main manuscript).

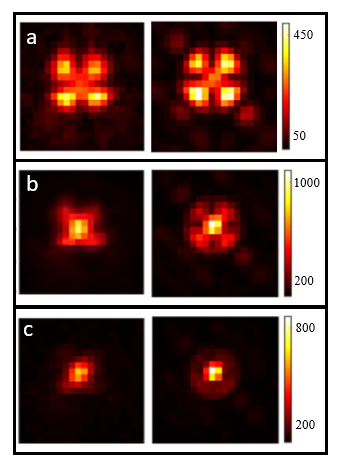

Fig. S 5 Data and Model for the XPSFs at focus. a) Red b) Orange c) Green XPSFs; Data on the left and the model on the right

Engineered PSFs often are largely dispersed in space with a large amount of the mass (intensity) of the PSF away from the center of the PSF. This may sometimes be a disadvantage when imaging multiple probes simultaneously, imaging a densely labeled sample, when there is a large background in the sample, when the photon budget is very limited, or when selecting the ROIs computationally. We obtained the radial sum of the XPSF and the Airy PSF to compare how the mass of the XPSF dispersed with the emission spectra and the depth. For this, we created the PSFs for our camera pixels and interpolated those PSFs over a grid of 100x100 pixels. Then, we obtained the sum of their mass in concentric circles starting from the center of the PSF at intervals corresponding to a single pixel (Orca Fusion-BT camera with 6.5-micron pixels) and obtained sums as shown in Fig. S 6 a) and b). We fitted a spline interpolant to the data points for continuity. Considering where 80% of the mass of the PSFs is contained, we calculate the dispersion (expansion) of the XPSF is only 30% (around 1 pixel radial expansion) when compared to a corresponding Airy PSF for the emitters near the focal plane and less than 1% for emitters away from the focal plane. We identify this as a positive trait of the XPSF when imaging multiple probes simultaneously as the level of overlap when compared to Airy PSFs is not significantly large. The sum at each sector is shown in Fig. S 6 c) and d). There is a difference of about 2 camera pixels between the segments where the largest PSF mass occurs between the Airy PSF and the XPSF near the focal plane and almost no difference when away from the focal plane.

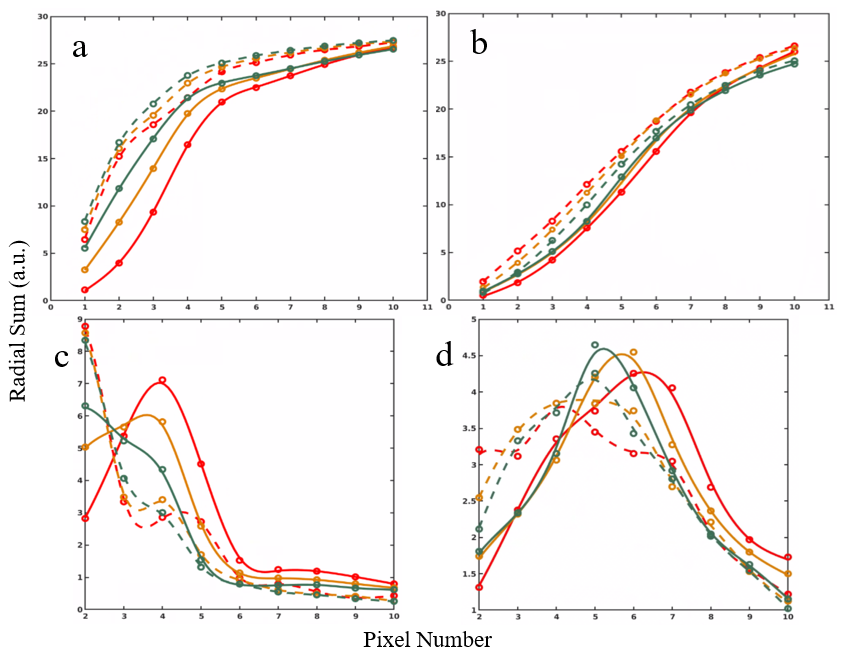

Fig. S 6 The intensity dispersion of the XPSFs compared to the Airy PSF. The cumulative radial sum of the intensity (mass) by considering increments of one pixel for the XPSF and the Airy PSF a) at z = 0 b) at z = 500 nm. The circular sector-wise sum c) at z = 0 d) at z = 500 nm. The solid lines correspond to the XPSF and the dashed lines to the Airy PSF. Red, orange, and green correspond to 680, 580, and 515 nm peak emissions for the three Tetraspeck bead spectra.

Fig. S 7 shows the truth table corresponding to the control experiment described in the
Discussion Section of the manuscript and Fig.S6b shows the truth table obtained through Monte-
Carlo simulations at around 100 photons per pixel background for a similar distribution of the
number of PSF photons in a sample. The truth table for the control experiment shows that the
PSFs belonging to AXF 488 have an increased chance of getting misidentified as that of CF 568 than
predicted by the simulations

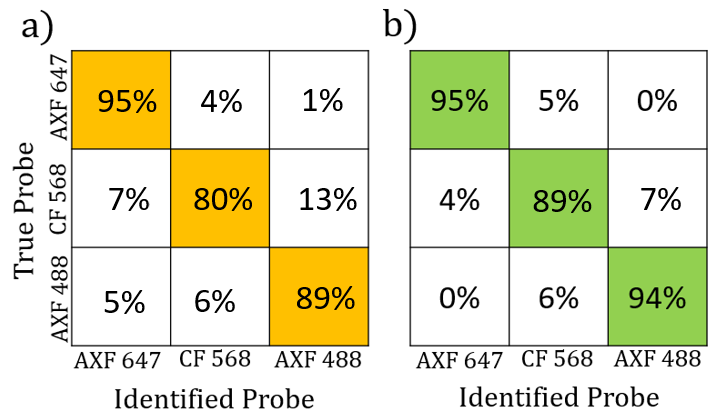

Fig. S 7 Truth tables describing the expected and the obtained spectral discernibility of the
XPSFs. The truth table corresponding to the observed results of the control experiment is in
panel a) and the results of the Monte-Carlo simulations for the best-case scenario described in
the Discussion Section of the main manuscript is in panel b). The background is around 100
photons per pixel and the mean number of PSF photons is around 2500 photons per PSF. The
diagonals in the truth tables correspond to the percentage of correct identifications for each
probe.

[1]	H. Kirshner, F. Aguet, D. Sage, and M. Unser, “3-D PSF fitting for fluorescence microscopy: implementation and localization application: 3-D PSF FITTING FOR FLUORESCENCE MICROSCOPY,” *J. Microsc.*, vol. 249, no. 1, pp. 13–25, Jan. 2013, doi: 10.1111/j.1365-2818.2012.03675.x.

[2]	M. Born and E. Wolf, *Principles of optics: electromagnetic theory of propagation, interference and diffraction of light*, 6th ed. Oxford ; New York: Pergamon Press, 1980. [Online]. Available: http://cdn.preterhuman.net/texts/science_and_technology/physics/Optics/Principles%20of%20Optics%20-%20M.Born,%20E.%20Wolf.pdf
